# WhichTF is dominant in your open chromatin data?

**DOI:** 10.1101/730200

**Authors:** Yosuke Tanigawa, Ethan S. Dyer, Gill Bejerano

## Abstract

We present WhichTF, a novel computational method to identify dominant transcription factors (TFs) from chromatin accessibility measurements. To rank TFs, WhichTF integrates high-confidence genome-wide computational prediction of TF binding sites based on evolutionary sequence conservation, putative gene-regulatory models, and ontology-based gene annotations. Applying WhichTF, we find that the identified dominant TFs have been implicated as functionally important in well-studied cell types, such as NF-κB family members in lymphocytes and GATA factors in cardiac tissue. To distinguish the transcriptional regulatory landscape in closely related samples, we devise a differential analysis framework and demonstrate its utility in lymphocyte, mesoderm developmental, and disease cells. We also find TFs known for stress response in multiple samples, suggesting routine experimental caveats that warrant careful consideration. WhichTF yields biological insight into known and novel molecular mechanisms of TF-mediated transcriptional regulation in diverse contexts, including human and mouse cell types, cell fate trajectories, and disease-associated tissues.

## Introduction

Transcription factors (TFs) are the master regulators of development. They define, refine, and can even divert cellular trajectories. TFs perform these important tasks by binding to specific DNA sequences in open chromatin, where they recruit additional co-factors and together modulate expression of downstream genes. TFs regulate biological processes in healthy adult tissues, and mutations to both TF genes and their genomic binding sites have been linked with human disease^1,2^.

The advent of next generation sequencing has paved the way for chromatin immunoprecipitation followed by sequencing (ChIP-seq)-based methods for the discovery of genome-wide loci where a given TF binds DNA in a given cell population^3^. Tools developed for the analysis of ChIP-seq data, such as GREAT^4^ (Gene Regulatory Enrichment of Annotations Tool), have discovered and leveraged a compelling phenomenon: when a TF is functionally important for the progression of a certain process, such that its perturbation leads to the disruption of this process, the binding sites for this TF are often highly enriched in the gene regulatory domains of the “downstream” target genes that drive this process^4^.

TFs work in different combinations to enact a vast repertoire of cellular fates and responses^5^. Between 1,500-2,000 TFs are thought to be encoded in the human genome1. Performing ChIP-seq for more than a handful of TFs in any cellular context is an expensive laborious procedure, while the assaying of hundreds of TFs even in the same cell state is impractical except in a handful of settings, by the most lavishly funded consortia.

To obtain a more comprehensive view of transcriptional regulation in action, experimental focus has turned from the assaying of individual TFs to the assaying of all open chromatin in a given cellular context. These DNase-seq, ATAC-seq, or single-cell ATAC-seq accessibility profiles offer a proxy for all cis-regulatory elements active in a given cellular state^6–8^.

While assaying all TFs is infeasible, many hundreds of TFs have been studied in one or more cellular contexts, or via complementary methods (such as protein binding microarrays or high-throughput SELEX), to obtain the DNA binding preference of the TF^1^. These hundreds of TF binding motifs can then be used to predict transcription factor binding sites (TFBSs) for all characterized TFs in various context-specific sets of accessible chromatin.

Very often, biological processes of interest are conserved at the genome sequence level across closely related species, such as primates or mammals. As such, computational tools like PRISM^9^ (Predicting Regulatory Information for Single Motifs) can be used to obtain a rarefied subset of binding site predictions that are both observed to be positioned in open chromatin and conserved orthologously in additional species. Because these sites evolve under purifying selection, they are more likely to be individually important in the probed context^9^.

Here, we innovate on the foundation of two tools our group previously developed: PRISM^9^ for the prediction of evolutionarily conserved binding sites for hundreds of human and mouse TFs, and GREAT^4^ for the detection of functions enriched in gene regulatory regions. We use insights from both to develop WhichTF, a tool that applies a novel statistical test to identify the most dominant TFs within a set of user-specified open chromatin regions. In this work, dominant TFs refer to TFs whose conserved binding sites are enriched within functionally-coherent regions of the input open chromatin regions. We show that our molecular definition of dominance successfully predicts biologically important factors in the context of different cell types, differentiation pathways, and even disease associated cellular sets.

## Results

### WhichTF Approach Overview

In order to predict dominant TFs, WhichTF relies on both functional genome annotations from GREAT and pre-curated, conservation-based predictions of TFBSs from PRISM. As such, we use GREAT in conjunction with the mouse genome informatics (MGI) phenotype ontology to annotate all genes in the human GRCh38 (hg38) and mouse GRCm38 (mm10) genomes with a canonical transcription start site (TSS), a putative gene regulatory domain, and any MGI phenotypes known to be affected by mutations to the associated gene. This procedure yields more than 700,000 gene-phenotype relationships for each genome (**Fig. 1a**, step 1)^4,10–12^. We also use PRISM to predict mammalian conserved TFBSs using 672 manually curated PWMs from 569 TFs across the entire genome^9^. The updated PRISM predictions resulted in 268 million and 161 million putative TFBSs for the human and mouse genomes, respectively (**Fig. 1a**, step 2).

**Fig. 1.**
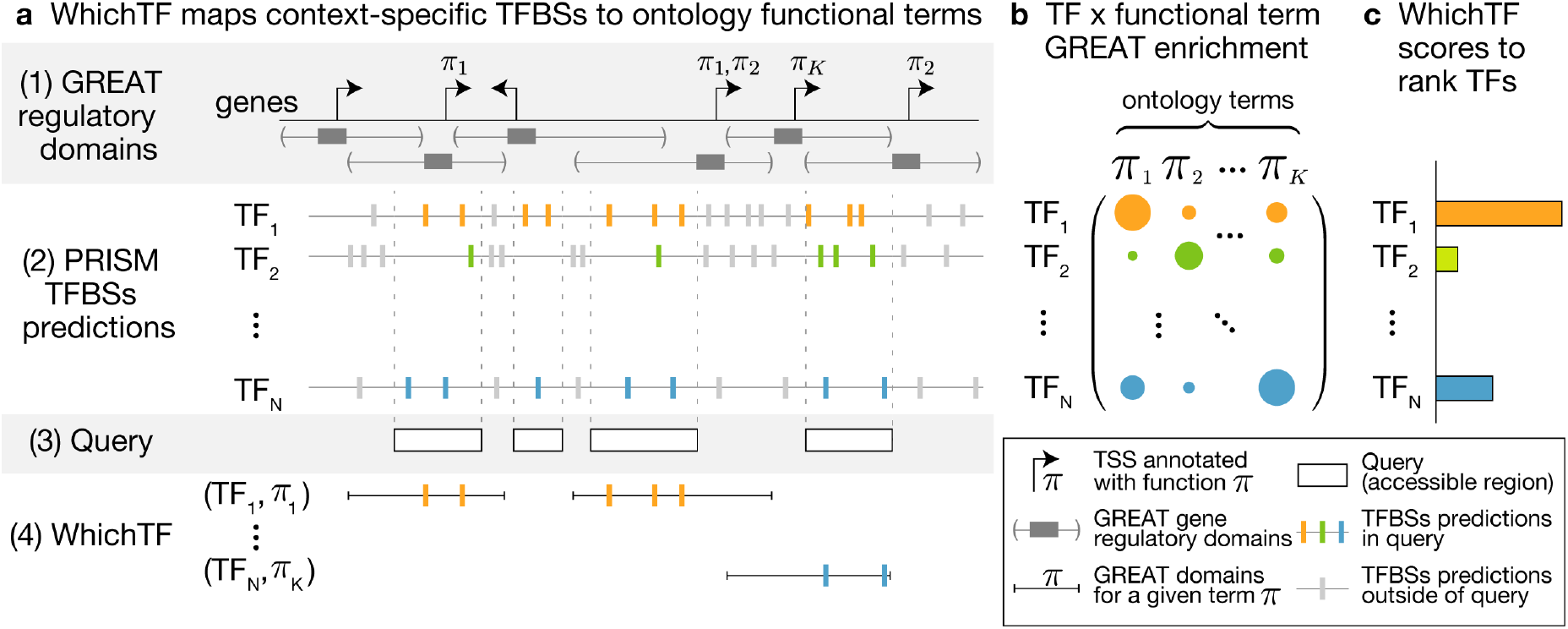
WhichTF identifies dominant TFs for given experimental measurements of chromatin accessibility. (a) WhichTF uses gene regulatory domain models and ontologies from the genomic region enrichment analysis tool (GREAT) (step 1) and conservation-based PRISM predictions of TFBSs (step 2). Given a user-defined set of genomic regions (step 3), WhichTF considers the top-*K* GREAT functional terms (*π*_1_, … *π*_*K*_) enriched in the query regions. For all pairwise combinations of top-*K* term and TF, WhichTF counts the number of TFBSs within the specified query regions (step 4). (b) The binomial and hypergeometric TFBS enrichment *p*-values for each ontology term are compiled in a TF-by-term summary statistic matrix. (c) Aggregating the summary statistics over terms, WhichTF returns a ranked list of TFs, ordered by predicted functional importance in the user-specific chromatin environment, with the corresponding scores and statistics (Online Methods). TSS, transcription start site.

To confirm the utility of restricting ourselves to regulatory domains of highly enriched ontology terms, we evaluated the relative enrichment in the number of TFBSs within the input open chromatin region as a baseline method (**Online Methods**). We found the baseline results are often overloaded with TFs associated with general housekeeping processes (**Supplementary Table S1**). We therefore turned to focus on the top 100 enriched terms (**Online Methods**).

For a given query (**Fig. 1a**, step 3), WhichTF uses functional annotations to enhance its prediction of dominant transcription factors. This is accomplished by computing TF enrichments in only a restricted, particularly relevant, subset of the user’s input. Specifically, WhichTF uses GREAT to identify enriched ontology terms within the user’s input query. Each term is associated with a region of the genome corresponding to all of the regulatory domains of genes annotated with that term. WhichTF selects the top 100 ontology terms. For each term and every TF, WhichTF counts the number of binding sites falling in the intersection of the user-specified accessible regions and the region of the genome associated to the term of interest (**Fig. 1a**, step 4), and computes enrichment statistics, represented as a TF-by-term enrichment matrix (**Fig. 1b**). Aggregating over the functional terms, WhichTF computes a novel score and significance used for ranking TFs (**Fig. 1c, Online Methods**). The top-ranked TFs are hypothesized to be functionally relevant TFs in a cell exhibiting the indicated accessibility profile.

### WhichTF identifies functionally important TFs across diverse cell types

To test the ability of WhichTF to identify functionally important TFs across different cell types, we applied WhichTF to DNase-seq profiles and found that the predicted dominant TFs are often confirmed to be functionally relevant by perturbation studies (**Fig. 2a**). In B- and T-cells, for example, we identified TFs in the NF-κB pathway, which are key factors in lymphocyte development and adaptive immunity^13^. In embryonic heart tissue, we found GATA-4, -5, and, -6 – known regulators of cardiac development and growth that, when perturbed, have been implicated in human congenital heart disease^14^. In embryonic hindbrain tissue, we found SOX2, a critical regulator of neural progenitor pluripotency and differentiation in embryogenesis and later development, including adult hippocampal neurogenesis^15–17^. WhichTF yielded similar biologically meaningful results from the corresponding cell types for mouse DNase-seq datasets (**Supplementary Table S2**), suggesting that WhichTF can highlight both the identity and evolutionarily conserved binding sites of key TFs from open chromatin in diverse contexts across species.

**Fig. 2.**
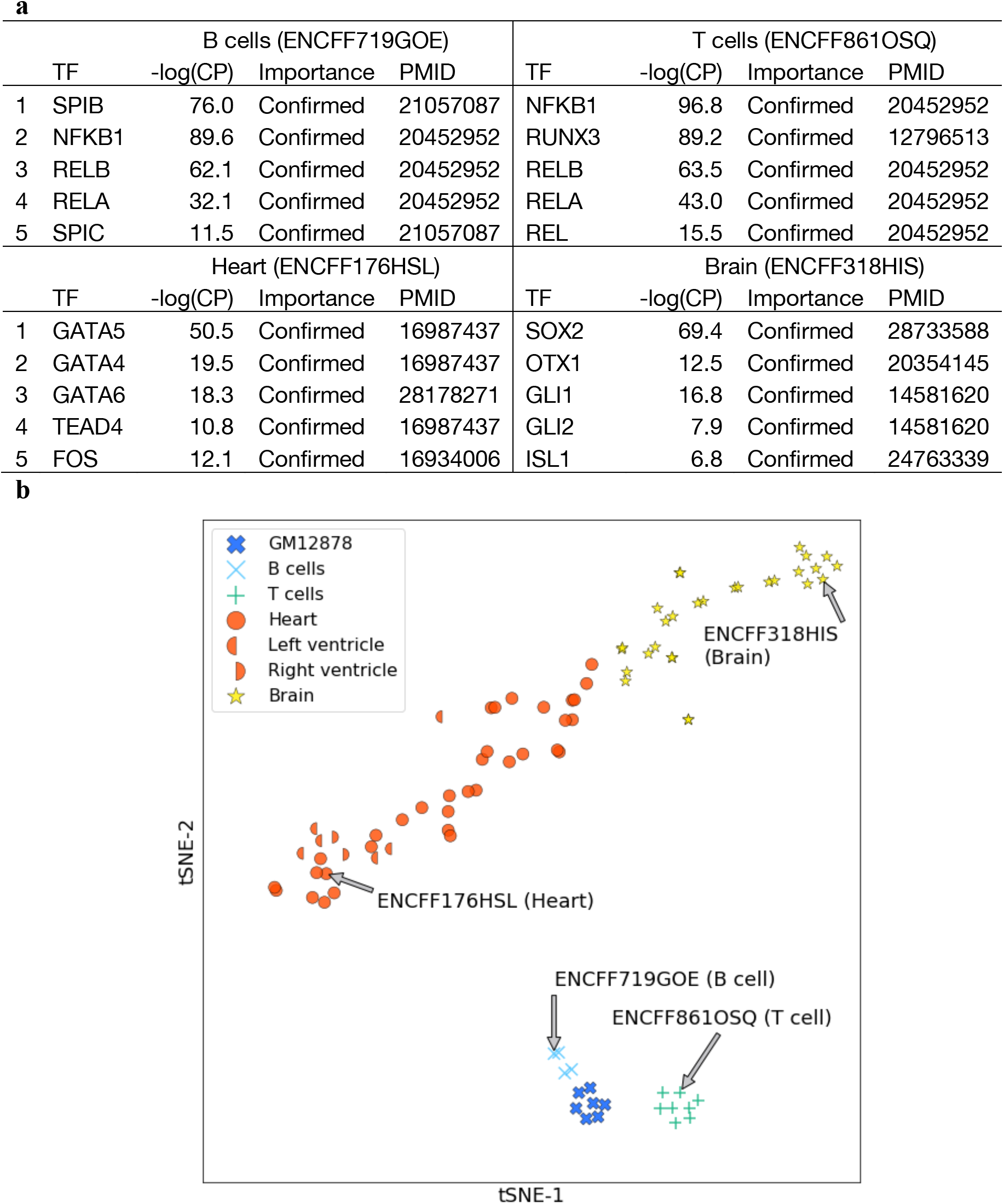
WhichTF identifies dominant TFs in diverse cell types. (a) The top 5 identified dominant TFs for B-, T-, heart, and brain cells are shown with the corresponding negative log conditional probability (-log CP), a statistical significance of the score of each TF, conditioned on the TFs with higher score (Online Methods). The importance and PubMed ID (PMID) columns indicate whether existing literature supports the role of the identified TFs, typically through perturbation experiments. (b) For DNase-seq data tracks of 90 samples across 7 cell types, the WhichTF score vectors are projected to t-SNE plot. WhichTF quantitatively and robustly captures biological similarities and dissimilarities of TF-mediated transcriptional programs. The samples highlighted in (a) are annotated with arrows.

### WhichTF robustly quantifies biologically meaningful similarities and differences in TF-mediated transcriptional programs

Precise knowledge of cell state and identity is crucial for understanding normal development and disease. To assess whether WhichTF can quantitatively and robustly capture biologically meaningful similarities and differences in TF-mediated transcriptional programs, we applied a t-distributed stochastic neighbor embedding (t-SNE) analysis to WhichTF score vectors computed for 90 samples across 7 cell types^18^. We found brain, lung, and hematopoietic cells are mapped to distinct regions (**Fig. 2b**). Furthermore, we saw fine-grained substructures among closely related samples. For example, we observed a clear separation of GM12878, B-cells, and T-cells. Reassuringly, different samples from the same biological tissue, such as left ventricle, right ventricle, and heart, showed no clear separation.

### WhichTF identifies differentially dominant TFs for closely related cell types

B-cells and T-cells share a closely related developmental trajectory^13^. As Fig. 2a shows, WhichTF identified NF-κB family members NFKB1, RELA, and RELB as shared dominant TFs. WhichTF also identified lineage-specific factors, such as SPI-B for B-cells and RUNX3 for T-cells (**Fig. 2a**). SPI-B is an ETS family TF known to play a key role in B-cell development and function, and environmental response^19–21^. RUNX3, in contrast, play T-cell-specific functional roles, such as in CD4 versus CD8 thymocyte commitment, helper versus killer T-cell specification, and helper type selection^22^. These differential roles for SPI-B and RUNX3 are corroborated by their cell-type-specific expression in B-cells and T-cells, respectively (**Fig. 3a**)^23^.

**Fig. 3.**
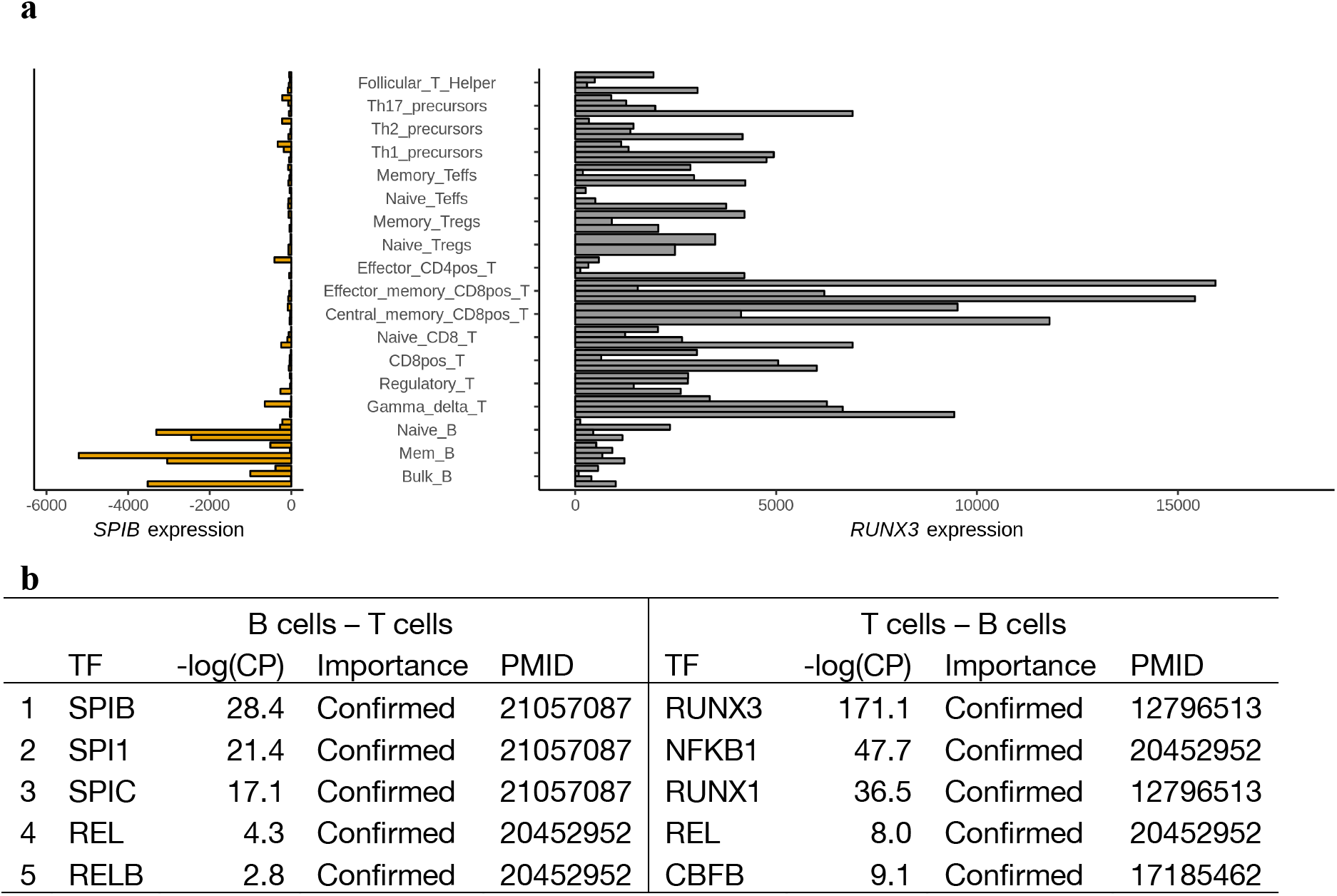
WhichTF identifies differentially dominant TFs in B and T-cell DNase-seq data. (a) Gene expression of the top differential TF genes, *SPI-B* and *RUNX3*, are shown (horizontal axis) across diverse lymphoid cell types (vertical axis) for up to four healthy donors. (b) The top 5 differential TFs for B-cells relative to T-cells (B-cell – T-cell) and vice versa (T-cell – B-cell) are shown with the corresponding statistical significance, negative log conditional probabilities (-log CP). The importance and PubMed ID (PMID) columns indicate whether existing literature supports the identified TFs.

Although we identified multiple TFs distinguishing B- and T-cells, the results are dominated by common factors. This is reasonable, as they share most of their developmental program^13^. To identify TFs with relative dominance from a given pair of samples, we developed a differential analysis framework focusing on uniquely accessible regions only in one sample (**Online Methods**). In B-cells, the differential analysis highlighted additional ETS family members, PU.1 and SPI-C. These TFs are essential for healthy B-cell differentiation and function (**Fig. 3b**). In T-cells, we saw an additional RUNX family member, RUNX1, as well as CBFβ (**Fig. 3b**) – both are functionally relevant in T-cells. Indeed, RUNX1, RUNX3 and CBFβ form a complex and are crucial for the healthy function of T-lymphocytes^32^.

### WhichTF identifies differentially dominant TFs along developmental trajectories

TFs regulate cell fate decisions in animal developmental programs^1^. To gain insights into the molecular mechanisms influencing cellular differentiation, we applied WhichTF to ATAC-seq data from timepoints along mesoderm development to identify differentially dominant TFs that distinguish cell fates at each step along the trajectory, from human embryonic stem cells (ESCs) to early somite vs. cardiac mesoderm (**Fig. 4**)^24^.

**Fig. 4.**
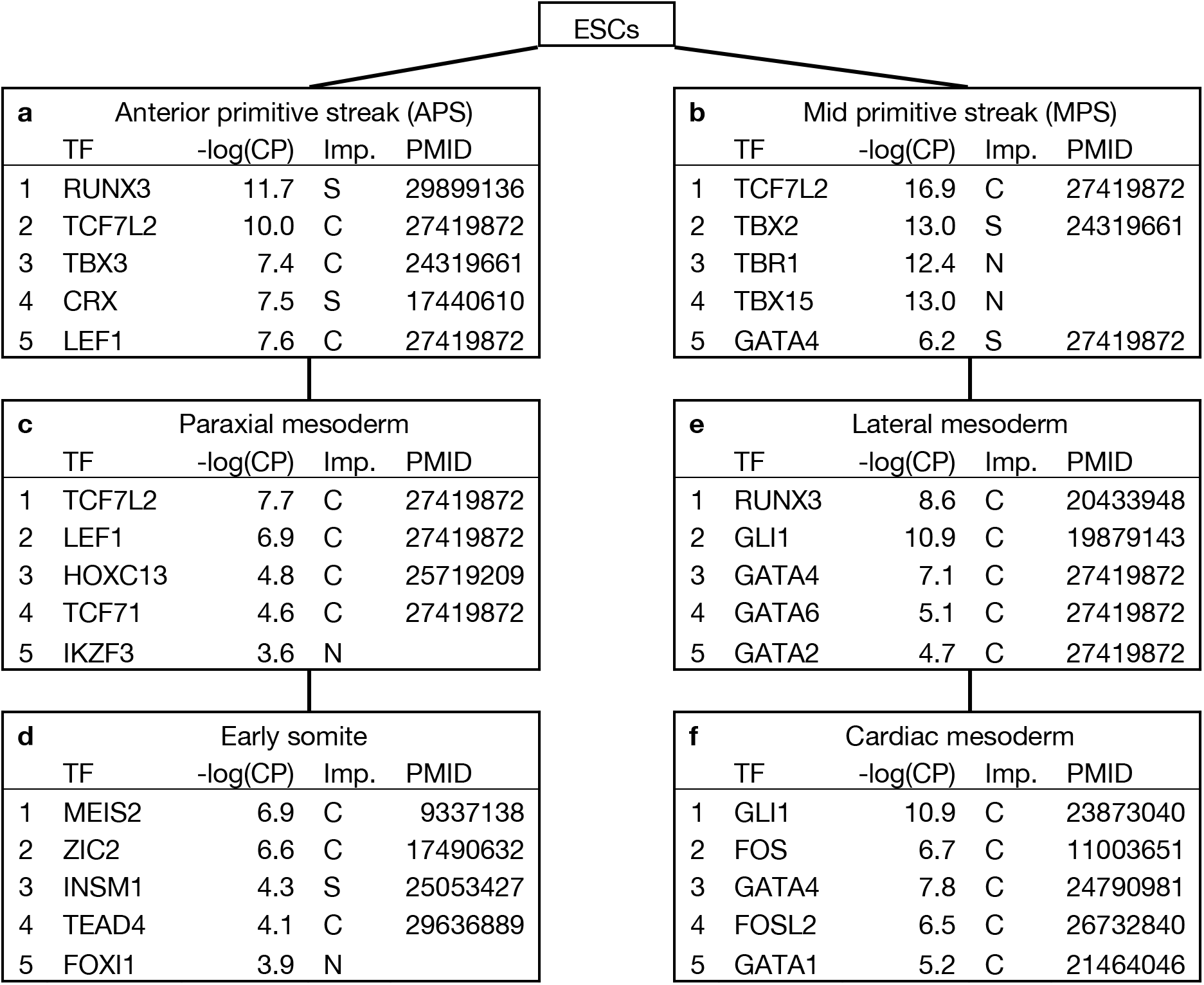
WhichTF identifies differentially dominant TFs compared to immediate progenitor cells along human mesoderm development pathway from ATAC-seq data. The top 5 TFs with the corresponding statistical significance, negative log conditional probabilities (-log CP) are shown. The importance (Imp.) and PubMed ID (PMID) columns indicate whether (i) existing literature supports the identified TFs (C: confirmed); (ii) literature reports closely related factors, such as co-factors and functionally related family members, or the identified TFs in related context (S: suggestive); or (iii) novel (N). ESCs, embryonic stem cells.

The first step of mesoderm development is the differentiation from ESCs to anterior (APS) or mid (MPS) primitive streak (PS) cells. In both APS and MPS cells, we found WNT signaling TFs, such as TCF7L2 and LEF1, as well as T-box family TFs, such as TBX-2 and -3 (**Fig. 4a-b**). WNT signaling is involved in PS differentiation and is crucial in inducing PS cell types^24^. T-box family members also play key roles in PS development. TBX6 is a canonical PS marker, and the specific loss of *Eomes* (a.k.a. *Tbr2*), causes ectopic primitive streak formation in mice^24,25^. The specific T-box family member TBX3, ranked third in APS cells, has been implicated in early stage of differentiation towards mesoderm from ESCs in mouse and *Xenopus* and has been reported for its functional redundancy with Tbx2 during *Xenopus* gastulation^26^. RUNX3, our top hit for APS, shows conserved expression in mouse neuromesodermal progenitor (NMP) cells and human D3-NMP-like cells. Interestingly, we also found previously unreported T-box family TFs, TBX15 and TBR1, of which TBX15 is linked to decreased skeletal muscle mass in mouse^12^ and known for tissue-specific expression in muscle, a tissue developed from the mesoderm lineage (**Supplementary Figure S1**).

In paraxial mesoderm, we found WNT signaling TFs, which promote paraxial and suppress lateral mesoderm (**Fig. 4c**)^24^. We also find HOXC13, necessary for proper development of the paraxial mesoderm into the presomatic mesoderm^27^. In early somites, we found MEIS2 and ZIC2, which are required in development of cranial and cardiac neural crest and somite cells, respectively (**Fig. 4d**)^28,29^.

In lateral mesoderm, we found multiple GATA family members, of which GATA4 is a downstream effector of BMP signaling in lateral mesoderm (**Fig. 4e**)^30^. We also saw *RUNX3*, which is co-expressed with *RUNX1* in lateral mesoderm^31^; both are necessary for hematopoiesis^22,32^. GLI1, a key TF in hedgehog (HH) signaling, is necessary for establishing left-right asymmetry in lateral mesoderm^33^. In cardiac mesoderm, we found FOS TFs, GATA TFs, and GLI1 (**Fig. 4f**). Interestingly, FOSL2 regulates the rate of myocardial differentiation^34^, and HH signaling via GLI1 is required for secondary heart field development^35^. As mentioned above, GATA factors are canonical drivers of cardiac development and all the GATA family members identified for mesoderm development (GATA-1, -2, -4, and -6) are implicated in Human cardiovascular diseases^14,2^.

### WhichTF identifies potentially disease-relevant TFs

Transcriptional mis-regulation has a broad impact on human diseases^2^. To assess whether WhichTF can shed light on the transcriptional regulatory molecular basis of human disorders, we examined systemic lupus erythematosus (SLE) as a case study. SLE is a heterogeneous and chronic autoimmune disorder most prevalent in young women and affecting 0.1% of the population. Its genetic and epi-genetic bases are poorly understood with known genetic associations accounting for only 10-20% of the observed heritability. While SLE is characterized by mis-regulated immune response in T- and B-cells, few TFs have been identified to play functionally relevant roles in SLE^36^.

To better understand the regulatory landscape of SLE, we identified differentially dominant TFs in healthy B-cells compared to SLE-affected B-cells and vice versa by applying WhichTF to ATAC-seq datasets^37^. We found BCL6 as a differentially dominant TF in healthy vs. SLE B-cells (**Table 1**). BCL6 is an important marker of T-helper follicular cells, a T-cell subtype which has been found to be mis-regulated in SLE^38^. Other differentially dominant TFs and their corresponding genes are implicated in autoimmune disorders (**Table 1**). A sonic hedgehog (SHH)-Gli signaling pathway member GLI1 is involved in pathogenesis of rheumatoid arthritis through synovial fibroblast proliferation^39^. A common genetic variant in *TCF7L2*, which is known for type 2 diabetes risk allele, discriminates autoimmune from non-autoimmune type 1 diabetes in young patients^40^. In a model system to study multiple sclerosis, ZEB1 is suggested as a regulator of experimental autoimmune encephalomyelitis^41^.

**Table 1:** WhichTF identifies disease relevant TFs. WhichTF identifies differentially dominant TFs from ATAC-seq measurement of B-cells from systemic lupus erythematosus (SLE) patients and healthy controls (HC). The top 5 TFs based on the analysis of HC with respect to SLE (HC - SLE) and vice versa (SLE HC) are shown with the corresponding statistical significance, negative log conditional probabilities (-log CP). The importance (Imp.) and PubMed ID (PMID) columns indicate whether literature supports the identified TFs: confirmed (C), suggestive (S), or novel (N).

| HC - SLE |  |  |  | SLE - HC |  |  |  |
| --- | --- | --- | --- | --- | --- | --- | --- |
| TF | -log(CP) | Imp. | PMID | TF | -log(CP) | Imp. | PMID |
| 1 BCL6 | 28.7 | C | 28045014 | GLI1 | 19.7 | S | 26552406 |
| 2 TFAP2B | 19.3 | N |  | ZFP143 | 11.0 | N |  |
| 3 ZEB1 | 16.6 | S | 20856809 | TCF7L2 | 6.0 | S | 18839133 |
| 4 ZSCAN21 | 15.2 | N |  | ONECUT2 | 5.2 | S | 28317889 |
| 5 ZSCAN20 | 14.2 | N |  | DMRTC2 | 3.8 | N |  |

### WhichTF uncovers stress response signatures

Context-specific measurements of open chromatin typically require purification of the desired cell type through mechanical and enzymatic tissue dissociation, which can be quite taxing on the cells. Indeed, it has been reported that stress response factors are often highly expressed in dissociated tissues^42^. Corroborating these observations, WhichTF often identifies canonical stress-associated TFs as some of the most dominant TFs in multiple very different contexts. As an illustration, we present WhichTF results for additional DNase-seq datasets (**Table 2**). For three endothelial cell types and adrenal gland cells, we found many members of FOS/AP-1 and NF-κB TFs, which are both known for their roles in stress response. We also found ZFP410 (also known as ZNF410), a poorly characterized Zinc finger TF, among the top hits across multiple cell types, suggesting its potential role in stress response. Even in the samples dominated by stress-associated TFs, we still found well-known context-specific players among the top hits, such as GATA3 and WT-1 in kidney cells and SOX and FOX TFs in endothelial cells^43–45^. We also found that the boundary between stress response and cell-type specific functions can be ambiguous, or at least context dependent. For example, we found FOS/AP-1 and NF-κB dominant in keratinocytes and B-cells, respectively which, in addition to being stress-associated, are also known for their context-specific functions^13,46^.

**Table 2:**
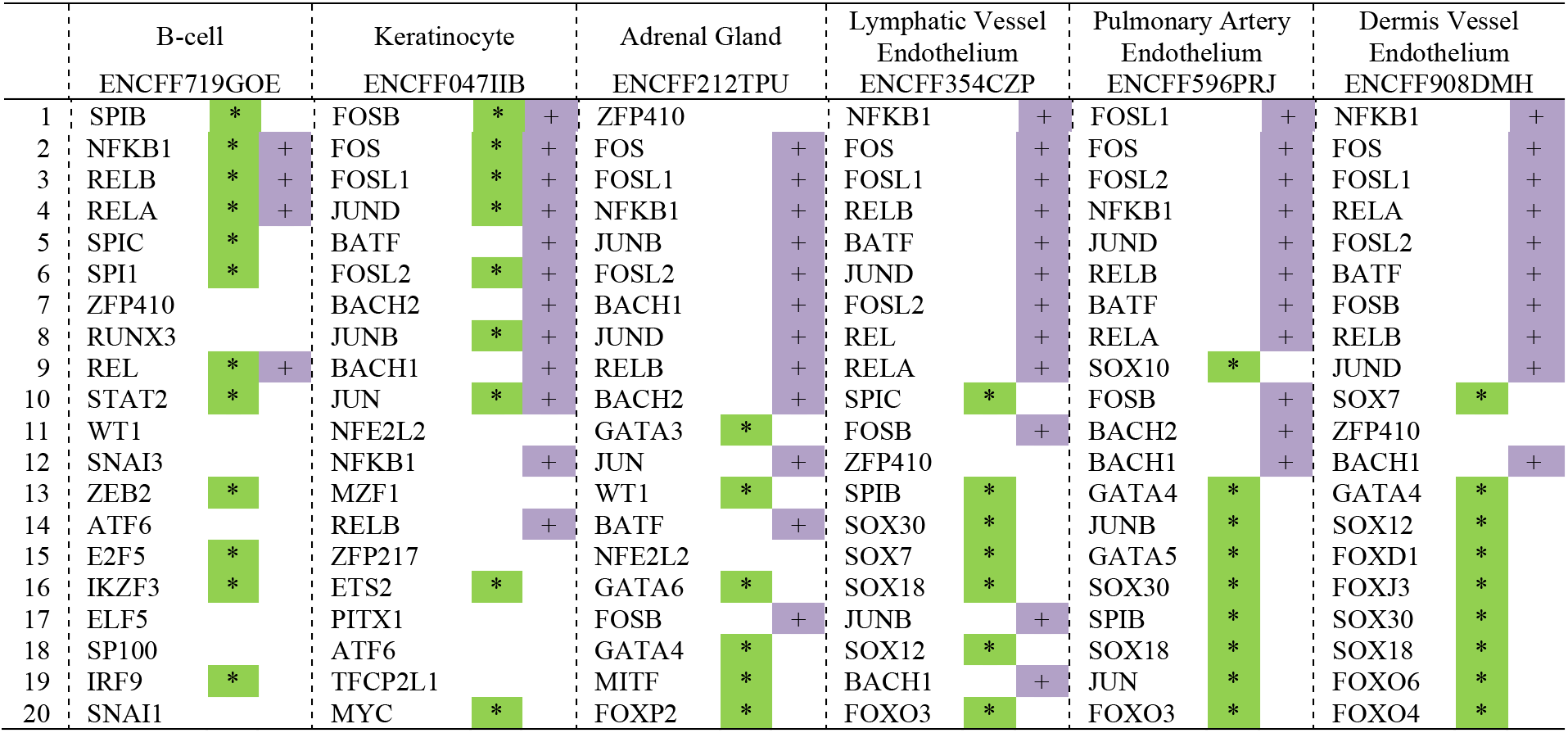
WhichTF identifies stress response factors in different samples. WhichTF identifies TFs known for stress response. The top 20 TFs identified by WhichTF are shown in ranked order for B-cells, keratinocytes, adrenal gland, lymphatic vessel endothelium, pulmonary artery endothelium, and dermis vessel endothelium cells. The TFs known to be involved in stress response signals are marked with plus (+), while TFs in families known to be functionally important in each context are marked with asterisk (*).

## Discussion

We present WhichTF, a novel computational method to identify and rank known or novel dominant TFs in any given set of accessible chromatin regions or through pairwise differential analysis of related samples. The WhichTF score is built on high confidence PRISM^9^ predictions of conserved TFBSs as well as gene regulatory domain and ontological annotation models from GREAT^4^. Applying WhichTF to dozens of samples across diverse biological contexts, such as multiple cell types, developmental programs, and disease samples, we found that the functional relevance of the identified dominant TFs is often supported or suggested by published literature.

WhichTF identifies not only cell-type specific TFs, but factors reflecting biological processes shared among multiple samples. One such example in our result, corroborated by previous expression profiling, suggests stress response due to cellular dissociation is a shared process^42^. In addition to previously identified factors, we report an under-characterized Zinc finger protein, ZNF410, as a TF potentially involved in cellular stress response. The identification of stress associated TFs suggests WhichTF may serve as a useful quality control of chromatin accessibility data.

As we have demonstrated above, WhichTF is broadly applicable. WhichTF takes as input any form of chromatin accessibility measurement for either human or mouse, the two most studied genomes. Our illustrative examples span both species and assay types, such as DNase-seq and ATAC-seq. When combined with emerging single-cell accessibility profiling technologies^8^, WhichTF will provide systematic characterization of dominant TFs across a spectrum of cell-types. For example, application of WhichTF to datasets from large-scale projects, such as the Human Cell Atlas project^47^, has the potential to discover dominant TFs for each cell type and binding sites of those TFs. Moreover, our differential analysis framework will help in understanding how closely related cell types diverge by providing hypotheses of differentially important TFs.

The resources made available with this study, including WhichTF and the GREAT update, provide an excellent foundation for investigating the molecular mechanisms of TF-mediated cis-regulation. Together, these results highlight the benefit of combining experimental characterization of chromatin accessibility, high-quality TFBS reference datasets, and ontological genome annotation, suggesting that systematic identification of dominant TFs across a large number of samples will be a powerful approach to understand molecular mechanisms of gene regulation and their influence on cell type differentiation, development, and disease.

## Online Methods

### GREAT v.4.0.4 update

We performed a major update of Genomic Regions Enrichment of Annotations Tool (GREAT)^4^ and released it as version 4.0.4. GREAT currently supports the human (*Homo sapiens* GRCh38 and GRCh37/hg19) and mouse (*Mus musculus* GRCm38/mm10 and NCBIM37/mm9) genomes. We obtained Ensembl gene sets from the following Ensembl^48^ versions:

- Human GRCh38: Ensembl version 90
- Human GRCh37: Ensembl for GRCh37 version 90
- Mouse GRCm38: Ensembl version 90
- Mouse NCBIM37: Ensembl version 67

By focusing on the set of genes with at least one Gene Ontology (GO) annotation^10,11^ as described before^4^, we defined putative gene regulatory domains for 18,777 (GRCh38), 18,549 (GRCh37/hg19), 21,395 (GRCm38/mm10), and 19,996 (NCBIM37/mm9) genes’ canonical transcription start sites.

We also updated the ontology reference data. GREAT currently supports the most recent versions of the following ontologies at the time of analysis: Ensembl genes, Gene Ontology (GO)^10,11^, human phenotype ontology^49^, and mouse genome informatics (MGI) phenotype ontology^12^ (**Supplementary Table S3**). The new Ensembl genes ontology is a “flat” ontology that makes every gene into a term, facilitating the testing of cis-regulatory elements congregation in the regulatory domains of individual genes. For MGI phenotype ontology, we mapped MGI gene identifiers to Ensembl human gene IDs using one-to-one orthology mappings from Ensembl Biomart^48^ version 90. In total, we compiled 2,861,656, 2,846,384, 2,734,172, and 2,675,691 gene-term relationships for GRCh38, GRCh37, GRCm38, and NCBIM37 genome assemblies, respectively (**Supplementary Table S3**).

### Computational TFBS prediction with PRISM

To take advantage of growing sequence data from both multiple species and functional genomics datasets, we updated our computationally predicted PRISM conserved transcription factor binding sites (TFBSs) for the human (*Homo sapiens* GRCh38 and GRCh37) and mouse (*Mus musculus* GRCm38 and NCBIM37) genomes. Briefly, PRISM predicts TFBSs based on evolutionary conservation of TF motif matches^9^. The GRCh37 and NCBIM37 tracks are derived using liftOver^50^ from that of GRCh38 and GRCm38, respectively.

We used the following multiple alignment from the UCSC genome browser^50^:

- Human GRCh38: Hg38 100-way conservation alignment (lastz)
- Mouse GRCm38: Mm10 60-way conservation alignment (lastz)

We removed Killer whale (*Orcinus orca*, orcOrc1) from the human alignment because of chromosome name mismatch. We further subset the alignments to Eutherian species^9^, resulting in 57 and 40 species for human and mouse, respectively. Using our manually curated TF monomer motif library^51^, we applied PRISM^9^ with the default parameters and focused on the top 10,000 predicted TFBSs for each TF in our analyses. We used GNU parallel in our analysis^52^.

### Baseline TF enrichment method without functional annotation

We computed the binomial p-value of each TFBS set, using the total number of TFBS predictions, the number intersecting the query and the fraction of the genome covered by the open chromatin region. We ranked the TFs by their binomial fold (**Supplementary Table S1**).

### WhichTF analysis protocol

WhichTF combines user specified accessibility measures, such as ATAC-seq or DNase-seq peaks with precomputed reference datasets to produce a ranked list of context specific, dominant TFs. The reference datasets consist of GREAT regulatory domain models, MGI mouse phenotype ontology-based gene annotations, and PRISM TFBS predictions.

WhichTF first identifies the top 100 ontology terms (*π*_1_, …, *π*_100_) based on the GREAT enrichment test on the input query set with the default “basal plus extension” association rule and a filter that terms must be associated with no fewer than two genes and no more than 500 genes associated to them. For each TF in the PRISM TFBS prediction library of *N* TFs, WhichTF takes an intersection of the TFBS prediction track and the user submitted open regions using overlapSelect^50^.

Each TF in the PRISM library has a different number of TFBSs and regulatory domains of different total sizes associated with each term. To capture the relative importance of different TFs within different contexts, WhichTF computes a few measures of statistical significance for each transcription factor and term and summarizes these measures in TF by term summary statistic matrices. Specifically, we apply hypergeometric and binomial tests defined below:

### TF hypergeometric test

Let’s define the GREAT gene regulatory domain for term *π*_*j*_ as RegDom_*j*_, PRISM TFBS prediction for TF_*i*_ as TFBS_*i*_, and user’s input query as QUERY. We define *n*_*i*_, *k*_*ij*_, *N*_*i*_, and *k*_*ij*_ as follows:

- *n*_*i*_ = #{TFBS_*i*_ ∩ QUERY}
- *k*_*ij*_ = #{(TFBS_*i*_ ∩ QUERY) ∩ RegDom_*j*_}
- *N* = #{(⋃_*k*_ TFBS_*k*_) ∩ QUERY}
- *K*_*j*_ = #{(⋃_*k*_ TFBS_*k*_) ∩ QUERY) ∩ RegDom_*j*_}

where, ∩ denotes genomic intersection operation and #{ *G* } denotes a function to count the number of elements in genomic regions, *G*. With these parameters, we compute the hypergeometric p-value for each pair of TF_*i*_ and term *π*_*j*_:

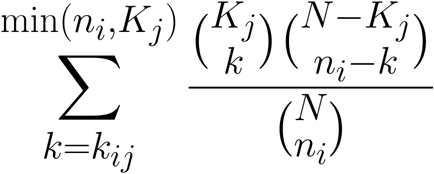

### TF binomial test

Using the intersection track, *TFBS*_*i*_ ∩ *QUERY*, we compute the GREAT binomial p-value for each pair of TF_*i*_ and term *π*_*j*_:

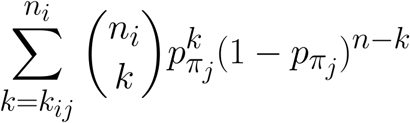

where, *p*_*π*_ denotes the probability of drawing a base annotated with term *π* from non-gap genomic sequences under the uniform distribution^4^.

### Adaptive TF significance threshold

To eliminate false positives, WhichTF focuses on terms where the most significant TF characterized by both hypergeometric and binomial p-value match. Using the enrichment statistics, WhichTF selects dominant TFs for each selected ontology term. We compute the adaptive threshold for each of the hypergeometric and binomial test by finding a leap in the p-values of the top 10 TFs for each term using the following procedure. Let’s denote the top 10 hypergeometric p-values for a fixed functional term *π* as *p*_1_ ≤ *p*_2_ ≤ … ≤ *p*_10_. We define the difference of adjacent negative log of p-values as 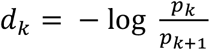. We define *m*, the index with the largest leap in p-value as *m* = argmax_*k*_ *d*_*k*_. Our adaptive threshold is *p*_m_ and we only keep TFs with hypergeometric p-values that satisfies *p* ≤ *p*_*m—*_. We define the adaptive threshold for binomial p-values in the same way. We say TF_*i*_ is significant for term *π*_*j*_ when it passes the adaptive thresholds for both TF hypergeometric and TF binomial tests.

### WhichTF scores

For each TF, WhichTF computes the score by the following equation. Let (*π*_1_, …, *π*_*K*_) be the set of terms selected from step 1 in the order of relevance with *π*_1_ as the top hit. Let Rank(TF_i_, *π*_*j*_) be the rank of the TF_i_ for term *π*_*j*_. Let Significant(TF_i_, *π*_*j*_G denote a Boolean variable that indicates whether TF_i_ passes the filters described above for term *π*_*j*_ (i.e. Significant is 1 if the TF passes the significance filter and zero otherwise). With this notation, we define the WhichTF score of TF_i_ as:

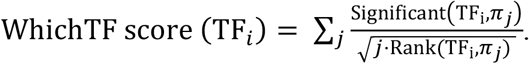

### WhichTF conditional p-values

WhichTF computes the statistical significance of a WhichTF score based on a null model that any ordering of TFs within each term is equally likely. Thus, the probability of a given score is determined by the relative number of configurations with the score. To enumerate the number of configurations with a given score in polynomial time, we devised a dynamic programing approach^53^ which acts recursively on the number of functional terms, *K*. This procedure first discretizes each contribution to the summand in the definition of the WhichTF score defined above. Let 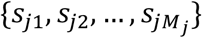 be the set of all the possible cumulative scores up to term *π*_*j*_, that is the scores gotten by computing the above sum only up to term *π*_*j*_. Here, *M*_*j*_ is the number of distinct discretized scores up to term *π*_*j*_. Let *n*_*ji*_ represent the number of different ways of getting each such score, *S*_*ji*_, and let 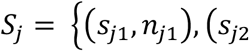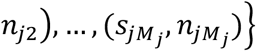 be the set of all tuples of scores and number of configurations. Finally, let 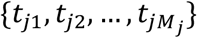 denote the individual summands at term *π*_*j*_.

The p-value of each score is computed directly from *S*_*K*_, the full set of cumulative scores and number of configurations, by dividing the number of configurations with scores greater than or equal to a given score by the total number of configurations. This list of tuples, *S*_*j*_, can be computed recursively with the base case of *S*_0_ = {(0, 1)}. The set of scores at level *j+1* is given by all combinations, *S*_*ji*_ + *t*_*j*+1*k*_, with the number of configurations given by aggregating over all combinations of *s* and *t* that yield the same cumulative score.

Given that the WhichTF scores of multiple TFs are not independent, we apply the procedure defined above from the top scoring TF to the TF with the lowest score and compute conditional statistical significance. This means that for the computation of statistical significance of the *i*-th ranking TF, we remove TFs whose rank is smaller than *i* and apply the recursive procedure defined above.

### Application of WhichTF in diverse functional contexts

#### Multiple cell types from the ENCODE/Roadmap project

From the ENCODE/Roadmap data portal, we obtained “hotspot” files derived from DNase-seq experiments^54,55^. All coordinates are provided in GRCh37. We present analysis spanning 95 samples from 12 cell types and tissues (Supplementary Table S4).

We systematically applied WhichTF to each sample and obtained the ranked list of TFs as well as a vector of WhichTF scores across all TFs in the library (**Figure 2a, Table 2**). We applied t-SNE, a non-linear dimension reduction method^18^, implemented in Python Scikit Learn library^56^ with perplexity 10 (**Figure 2b**).

Using mouse ENCODE DNase-seq datasets provided in GRCm38 from the four cell types used for the human analysis (**Figure 2a, Supplementary Table S5**), we applied WhichTF using mouse GRCm38 reference dataset (**Supplementary Table S2**).

#### Cell type-specific expression analysis

We presented cell type-specific RNA-seq data from the GEO database (GSE118165)^23^. We subseted this dataset to the unstimulated samples and plotted the expression of *SPIB* and *RUNX3* for lymphoid cells in T and B cell lineages (**Figure 3a**).

#### WhichTF for differential analysis

To find TFs dominant in an input set A compared to another input set B, we defined set A and set B regions as foreground and background, respectively. We used bedtools^57^ “subtract” to keep a subset of A that does not overlap with B. We applied WhichTF single run mode (above) on the identified differentially accessible regions (**Figure 3b**).

#### Mesoderm lineage dataset

Using ATAC-seq datasets (SRP073808 from NCBI GEO database) of mesoderm development^24^ (**Supplementary Table S6**), we applied WhichTF differential analysis following the diagram of sequential differentiation (**Figure 4**).

#### Systemic lupus erythematosus dataset

Eight sets (4 SLE and 4 healthy controls [HC]) were taken from the NCBI sequence read archive (SRA, **Supplementary Table S7**). Paired end reads were mapped using bowtie2 with the outer distance flag (-X) set to 1000 and otherwise default settings^58^. Samtools was used to generate a sorted bam file and MACS2 was used to call peaks with shift set to 37, extension size set to 72 and broad and keep-dup flags on^59,60^. Given that some of the samples in this dataset are from a biobank, we conservatively defined differentially accessible regions shown below and applied WhichTF differential analysis (**Table 1**):

- SLE − HC := SRR3158183 − ⋃_*x* ∈ SRR3158176-9_ *x*
- HC − SLE := ⋂_*x* ∈ SRR3158176-9_ *x* − ⋃_*x* ∈ SRR3158180-3_ *x*

#### Tissue-specific gene expression of the identified TF

Using the data obtained from the GTEx Portal^61^ on 05/24/2019 (phs000424.v7.p2), we investigated whether the identified TFs in have a tissue-specific expression (**Supplementary Figure S1**).

## Data availability

All datasets analyzed in this study are publicly available through the ENCODE/Roadmap portal [https://www.encodeproject.org/], NCBI GEO database [https://www.ncbi.nlm.nih.gov/geo/], NCBI sequence read archive [NCBI sequence read archive], or the GTEx Portal [https://gtexportal.org] with identifiers included in Supplementary Tables S4-S7 and in Online Methods.

## Code availability

WhichTF program and analysis scripts are available at our Bitbucket repository: https://bitbucket.org/bejerano/whichtf

GREAT version 4.0.4: https://great.stanford.edu

## Acknowledgements

We thank Stanford’s Kyle M. Loh, as well as Heidi Chen, Alex M. Tseng, and other members of the Bejerano Lab for useful discussions, feedback and advice. Y.T. is supported by a Funai Overseas Scholarship from the Funai Foundation for Information Technology and the Stanford University School of Medicine. E.S.D. was supported in part by the Simons Collaboration Grant on the Non-Perturbative Bootstrap. This work was supported by National Institute of Mental Health (NIMH) of the National Institutes of Health (NIH) under awards U01MH105949 to G.B. The content is solely the responsibility of the authors and does not necessarily represent the official views of the National Institutes of Health.

## Author information

### Author contributions

E.S.D., Y.T. and G.B. conceived and designed the study. Y.T. updated GREAT. E.S.D. conceived of and developed the WhichTF algorithm with support from Y.T. and G.B. E.S.D. and Y.T. performed the computational analyses. Y.T. led the completion of the manuscript with support from E.S.D. and oversight from G.B. Y.T. and E.S.D. contributed equally to the project and author list is ordered by age. The manuscript was written and approved by all authors.

## Supplementary materials

### List of supplementary materials

#### Supplementary Figures

- Supplementary Figure S1: Gene expression profile of *TBX15*

#### Supplementary tables

- Supplementary Table S1: Baseline TF enrichment method
- Supplementary Table S2: Mouse ENCODE dataset analysis
- Supplementary Table S3: The update summary of GREAT ontologies
- Supplementary Table S4: Human ENCODE datasets
- Supplementary Table S5: Mouse ENCODE datasets
- Supplementary Table S6: Mesoderm development samples
- Supplementary Table S7: Sequence read archive accession IDs for systemic lupus erythematosus dataset

#### Supplementary Figures

**Figure S1.**
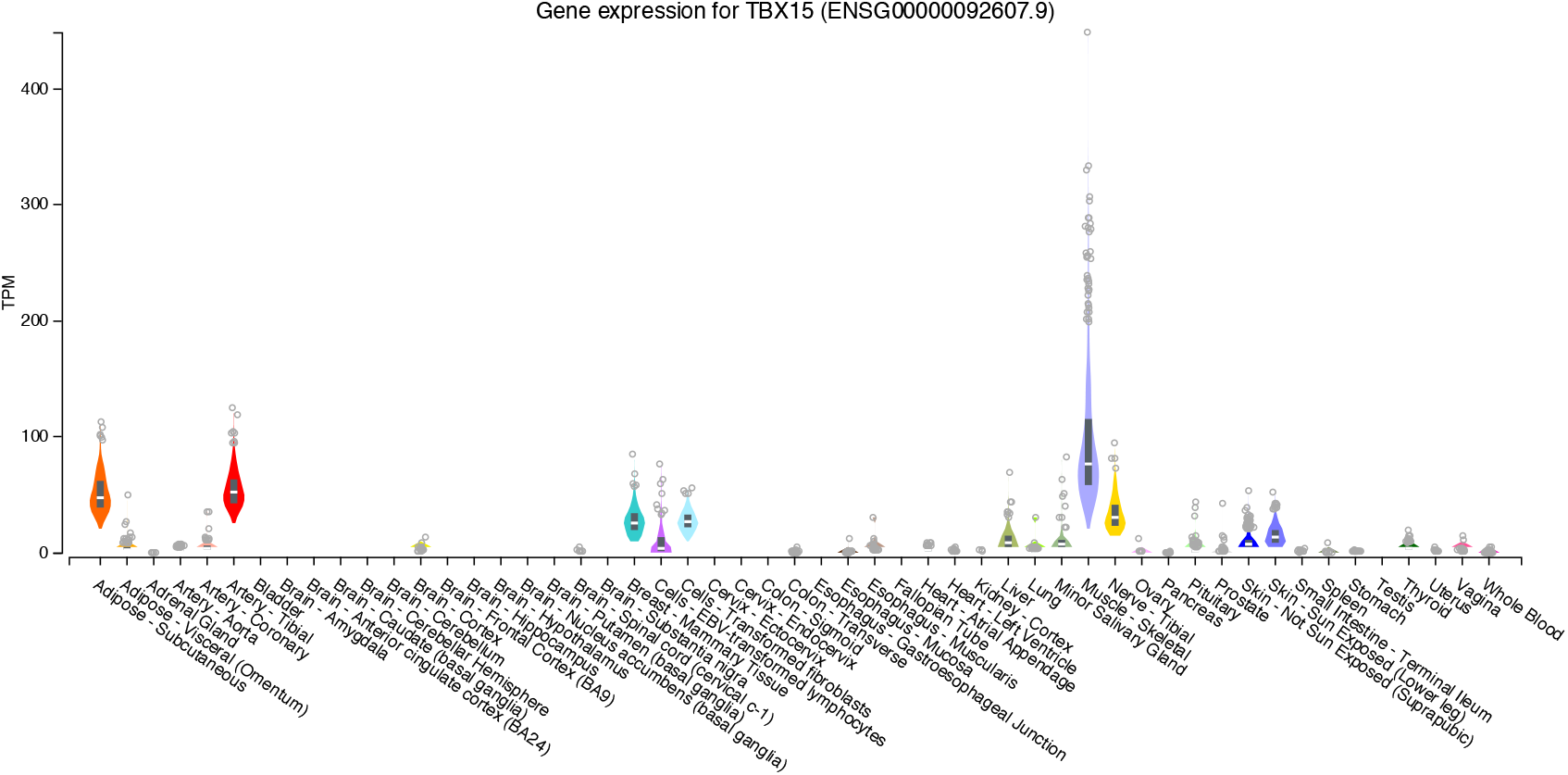
Tissue-specific gene expression profile of *TBX15* in muscle. The Human cell types are shown on x-axis and the expression (TPM) is shown on y-axis. The median and 25th and 75th percentiles are shown as box plots and data points are shown as outliers if they are above or below 1.5 times the interquartile range.

#### Supplementary Tables

**Supplementary Table S1.** Baseline TF enrichment method for the four human cell types from ENCODE and Roadmap DNase-seq datasets are shown. The top 5 identified TFs are shown for (a) B-cells, (b) T-cells, (c) heart cells, and (d) brain cells. ENCODE accession IDs for each sample and the dominant TFs and their corresponding −log10(p-value) are shown. There is less cell-type specificity in the identified results.

**Supplementary Table S2.** Mouse ENCODE dataset analysis. WhichTF identifies dominant TFs for four mouse cell types from ENCODE and Roadmap DNase-seq dataset. The top 5 identified dominant TFs are shown for (a) B-cells, (b) T-cells, (c) heart cells, and (d) hindbrain cells. The ENCODE accession IDs for each sample are shown on the top and the dominant TFs and their corresponding statistical significance, conditional probabilities, are shown.

**Supplementary table S3.** The update summary of GREAT ontologies. Ensembl genes is a flat ontology defined from the set of genes with at least one meaningful annotation in gene ontology (Online Methods). GO: gene ontology. HPO: human phenotype ontology. MGI: mouse genome informatics.

**Supplementary Table S4.** Human ENCODE datasets. The list of ENCODE accession IDs used in our study and the corresponding cell type or tissues.

**Supplementary Table S5.** Mouse ENCODE datasets. The list of ENCODE accession IDs used in our study and the corresponding cell type or tissues.

**Supplementary Table S6.** Mesoderm development samples. The list of sample IDs, sample description, and the reference to the corresponding results.

**Supplementary Table S7.** Sequence read archive (SRA) accession IDs for systemic lupus erythematosus dataset. SLE indicates disease and HC indicates healthy control.

